# How important is the intra-regional soil heterogeneity for the design of future stress-avoidant wheat ideotypes? A modeling study in central France

**DOI:** 10.64898/2026.02.11.705307

**Authors:** Guillaume Blanchet, Mikhail A. Semenov, Vincent Allard

**Author notes:** Corresponding author: Vincent Allard.

## Abstract

Accurate projections of crop adaptation to climate change require accounting for the spatial heterogeneity of soils, which modulates both water availability and the effectiveness of genetic adaptation. Using the process-based crop model Sirius, we investigated how intra-regional variability in soil available water capacity (AWC) influences wheat yields and the adaptive value of stress-avoidant ideotypes under future climates in central France (Limagne plain). Detailed soil databases were aggregated across five representative sites and combined with multiple climate projections (CMIP6), two emission pathways (SSP2-4.5 and SSP5-8.5), and three time horizons (2031–2050, 2051-2070 and 2071-2090).

Variance decomposition revealed that soil AWC accounted for 23% of the simulated yield variability, significantly exceeding the contribution of local climate contrasts (10%), a pattern consistent across current and future periods. Deep soils (>80 mm AWC) buffered drought effects whereas yields stagnated in shallow soils (<80 mm AWC) where water deficits persisted despite phenology hastening. On average, the reference cultivar showed earlier anthesis by 8–21 days under future climates, leading to higher yields mainly in deep soils. Optimization of flowering timing through stress-avoidant ideotypes provided mean yield gains of +6.33 dt·ha^−1^ in deep soils, but limited benefits (+1.71 dt.ha^-1^) in shallow ones, highlighting pedological dependence of breeding efficiency.

Advancing anthesis also increased exposure to early-spring frost: frost probability rose from <0.1 to >0.4 when flowering occurred more than 250 °C.days earlier, particularly in the frost-prone part of the study area. Hence, frost risk remains a critical constraint for early ideotypes, even under strong warming.

Overall, our results demonstrate that intra-regional soil heterogeneity remains a dominant driver of wheat yield variability and adaptation potential under climate change. Designing stress-avoidant ideotypes without explicit consideration of local soil AWC could lead to maladaptation, especially in regions with shallow soils represent a significant portion of cropped areas. In such situation, breeding for terminal stress avoidance may offer only limited benefit. We advocate that breeding and modeling frameworks integrate high-resolution soil data to refine regional ideotype design, reconcile terminal-stress avoidance with frost tolerance, and better capture the spatial realism required for sustainable crop adaptation strategies.

**Highlights:**

- Local soil water capacity limits wheat adaptation to climate change.
- Deep soils favor earlier, stress-avoidant ideotypes.
- Shallow soils restrict the benefits of phenological adjustment for stress avoidance.
- Frost exposure remains a key risk when shifting phenology toward earliness.

## Introduction

One of the most significant challenges to be overcome in the context of climate change is the regional adaptation of agricultural practices. Under the anthropogenic increase of greenhouse gases concentration in the atmosphere, climate changes are already observed or expected in the near future. Mean surface temperature increases globally, but with strong regional and seasonal disparities. For example, western Europe is currently experiencing an extremely fast warming that is twice the rate observed globally (WMO, 2023), particularly in spring and summer (Ribes et al., 2022; Schumacher et al., 2024). The likelihood of extreme temperature events, such as summer heat waves, has already increased dramatically, with heat events expected twice a century in the early 2000s now expected to occur twice a decade (Christidis et al., 2015). Both aspects, trend and hazards likelihood, are expected to be amplified in the near future (IPCC, 2023). Similarly, precipitation patterns are also altered, with disparate trends observed at the regional and continental scales. Modifications of rainfall regime are already observed in Europe, where annual precipitation increased in northern Europe over the last half century, while a gradual decline is occurring in the Mediterranean basin (Spinoni et al., 2017; Stagge et al., 2017). Extreme precipitation events, such as severe droughts and heavy precipitation, are also becoming more frequent and longer lasting (Grillakis, 2019). In Western Europe, winter wheat is a major staple crop and its production is already facing the increasing impacts of climate hazards (Brisson et al., 2010). Among the consequences, yield stagnation is observed in spite of the ongoing genetic improvements achieved in the last decades (Le Gouis et al., 2020; Oury et al., 2012).

Several agricultural adaptation strategies can be implemented to achieve sustainable production in relation to climate change. A large number of these strategies rely on modifications at the farm level (*e*.*g*., diversification of production) or at the plot level (e.g., adaptation of sowing date, tillage practices, …). Other options rely on breeding cultivars specifically adapted to the projected climate conditions. Two main approaches have emerged for adapting wheat. One is to minimize exposure to climate hazards during sensitive phenological stages, *i*.*e*., stress avoidance. The alternative strategy involves enhancing crop tolerance to projected climate conditions, with substantial investment directed toward genetic improvement of stress tolerance. A large number of genes or quantitative trait loci (QTLs) have been identified with respect to drought and heat tolerance (Pinto et al., 2010). Nevertheless, these works did not result in a significant improvement in yield under stressful conditions, due to the highly polygenic nature of the related traits (Fleury et al., 2010). In addition, the adaptive traits (and the underlying genes) conferring some forms of tolerance are highly dependent on the exact nature of the stress occurrence (Tardieu, 2012) and thus hardly usable in real, changeable agronomic conditions. Conversely, strategies that rely on stress avoidance may appear more pragmatic. Indeed, the majority of genes related to wheat phenology have already been identified (Hyles et al., 2020) and the associated genetic variability is sufficiently large to design wheat ideotypes that are extremely contrasted in terms of phenology (Celestina et al., 2023). This approach has already been extensively explored in order to identify wheat ideotypes adapted to specific climate scenarios (Rötter et al., 2015; Semenov & Stratonovitch, 2013, 2015). One of the key issues is the existing trade-off between terminal stress avoidance, which favor earlier genotypes, and biomass accumulation, which is generally enhanced in case of late genotypes (Asseng et al., 2015). Additionally, the exposure of early genotypes to frost risk should not be overlooked as they might still represent a critical hazard for crops in the near future (Zheng et al., 2012, 2015). Consequently, the use of modelling studies to accurately frame the appropriate phenology for these ideotypes is necessary. In this regard, crop models represent valuable tools since they allow to integrate future climate scenarios, soil characteristics, and technical management in order to predict crop performance.

Extensive research has been conducted to model the wheat phenology based on various factors, including the influence of temperature (Jamieson et al., 1995; Kirby, 1990), vernalization requirements (Robertson et al., 1996), and photoperiod sensitivity (Brooking et al., 1995; Brooking & Jamieson, 2002). In the continuity of these efforts, studies showed that it is possible to link phenological parameters with identified genes or QTLs (Bogard et al., 2014, 2020; He et al., 2012), opening the way to the virtual exploration of the current germplasm and its suitability to future conditions (Zheng et al., 2016). So far, the emphasis has been put on thermal stresses and simulations studies suggest that early heading cultivar would be efficient to reduce the exposition to terminal heat (Gouache et al., 2012), but late frost should not be overlooked because it might still represent adverse conditions in specific wheat-growing regions (Bogard et al., 2021; Zheng et al., 2012). However, the scope of crop models’ applicability remains constrained by the simplistic (or nonexistent) representation of specific processes or the scale of the study. For instance, water stress has not been considered to identify optimal phenology under future climate scenarios, though its relevance has been already assessed (Flohr et al., 2017).

Prospective studies for future climate scenarios have mostly been conducted at relatively large scales, where both climate projections and soil conditions are highly contrasted. At this level of aggregation, the effects of climate variability are often emphasized, while the role of soil heterogeneity remains less explored (Waha et al., 2015). However, recent findings indicate that uncertainties in soil data can rival or even exceed the effects of climate signals on yield projections, particularly under low-input conditions (Folberth et al., 2016). At the intra-regional scale, soil heterogeneity also emerges as a major driver of yield variability, although its relative importance compared with climate variability depends on the context (Wassenaar et al., 1999). Therefore, recommendations derived from large-scale simulation studies may not be adequate for guiding varietal choices at finer spatial scales.

We propose to evaluate the significance of soil heterogeneity at the intra-regional scale, with regards to stress-avoidant wheat ideotypes designed for different climate scenarios. To this end, we performed a modeling study of designing wheat ideotypes over a spatially limited territory in central France, the Limagne Basin, where detailed soil information was available. First, we analyzed the hierarchy of pedoclimatic factors on simulated wheat yields at the scale of the Limagne territory according to crop models. We subsequently assessed the implications of climate change effects on simulated phenology and yield on a current reference genotype. Finally, we evaluated the adaptation potential of a reference wheat genotype in the Limagne by optimizing only the date of anthesis and we characterized possible adaptation with regards to local soil conditions.

## Materials and Methods

### Sites and soil data

The Limagne Basin is an agricultural plain situated in central France, in the vicinity of Clermont-Ferrand (Fig. 1). The plain covers approximately 200,000 ha and is mainly cropped with wheat, grain maize and sunflower. Local wheat production is of significant economic importance and largely depends on high-input practices. The plain is a north-south oriented half-graben that has been filled with detrital materials, sediments and alluviums over successive geological cycles (Vennin et al., 2021). Local pedogenesis has resulted in a variety of soil types, ranging from shallow to deep calcareous clay soils (calcisols) to heavy and deep clay soils (vertisols). In the eastern part of the plain, the Allier River formed sandy alluvial soils (fluvisols) (Nowak & Marliac, 2020). The north-south extension of the plain covers approximately 150 km, with varying degrees of orographic protection, resulting in local climatic contrasts (see the next subsection for detailed information). To represent the local pedoclimatic gradient and the associated soil heterogeneity, five sites were selected along the Limagne basin for analysis: Montbeugny (MB), Vichy (VC), Clermont-Ferrand (CF), Issoire (IS), and Fontannes (FT) (Fig. 1).

**Fig. 1.**
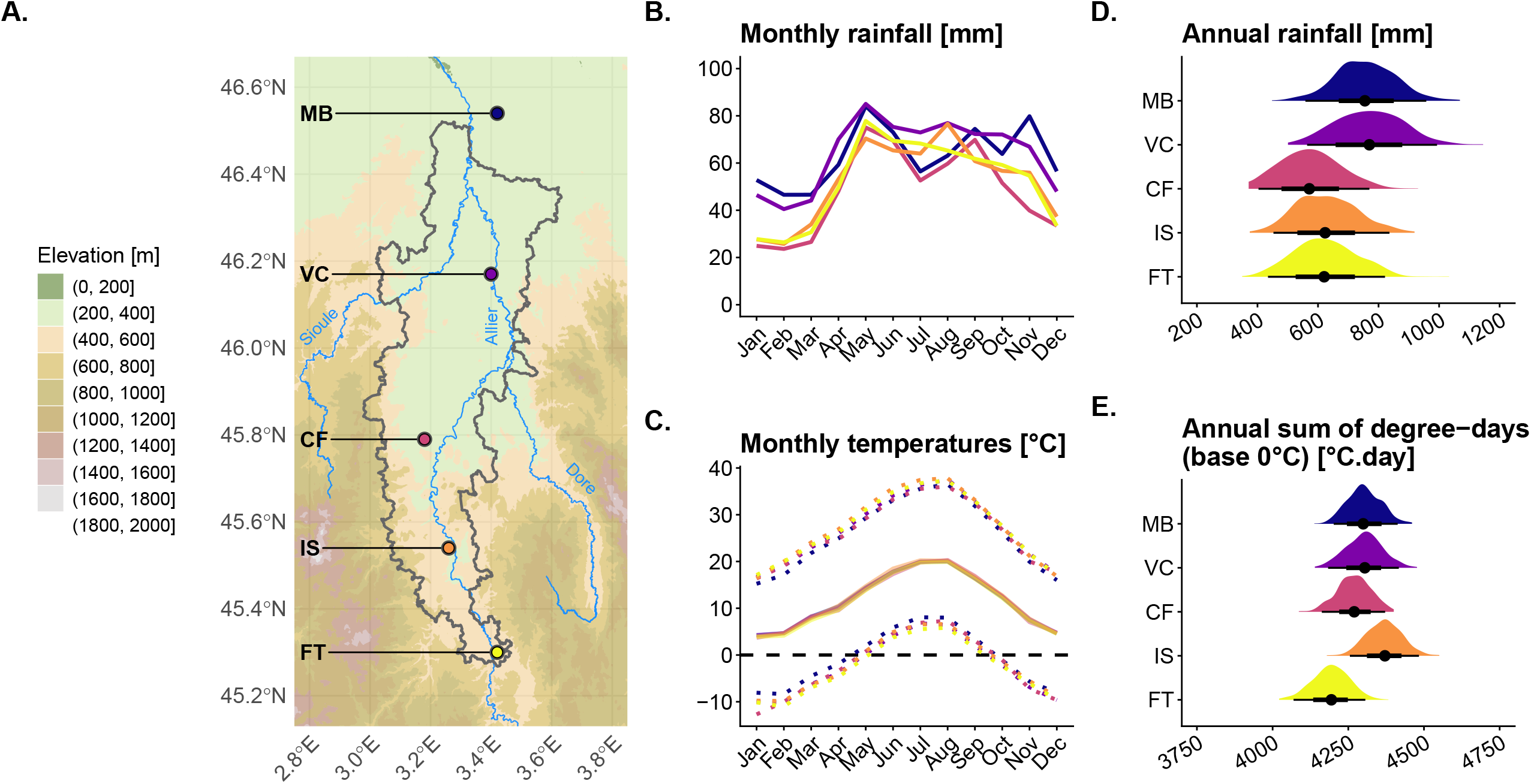
Rainfall and thermal regimes of the five study sites in the Limagne plain over the reference period (1991-2020 ; weather dataset generated with LARS-WG). The map’s color scale shows the topography, while the thick gray line delineates the regional boundaries of Limagne, as proposed by the French National Institute for Statistical and Economic Studies (INSEE) for classifying French agricultural regions. Solid lines depict mean monthly statistics (B. and C.), while dotted lines depict average minimal and maximal temperatures (C.). Frequency plots (D. and E.) display the distribution of respective variables over the 500-year generated dataset.

Regional soil information was obtained by accessing and aggregating three existing soil spatial databases established at a scale of 1:250,000 (Bornand et al., 1966; Genevois et al., 2019; Pelletier et al., 2012). In these databases, soil information is structured into fully georeferenced soil mapping units (SMU), and further decomposed into non-explicitly georeferenced soil topological units (STU). For each STU, a detailed description of individual soil layers (*e*.*g*. soil texture, soil organic matter content, stone content, layer’s thickness, etc.) is provided, as well as a percentage cover of the SMU. Pedological information of arable land was synthesized by filtering SMU covered by field crops only. To that purpose, areas cultivated either with winter or spring crops were extracted from the 2014-2021 graphic parcel register, *i*.*e*. a georeferenced database established according to farmers’ declarations within the context of the Common Agricultural Policy (available at https://www.data.gouv.fr/fr/datasets/registre-parcellaire-graphique-rpg-contours-des-parcelles-et-ilots-culturaux-et-leur-groupe-de-cultures-majoritaire/).

The soil information was synthesized in a stepwise manner to ensure the quality of the information on soil available water capacity (AWC) over the spatial scales considered, while making compromises to simplify its virtual representation with the crop model used in the present study, *i*.*e*., Sirius (Jamieson et al., 1998). First, all soil profiles from every considered STU were simplified as a single homogeneous soil layer. Based on local expert knowledge, only root colonizable soil layers were considered within the soil profile, thus limiting soil depth (Δz). Then, soil texture and stone content (□_stone_) were averaged over the entire soil profile, weighted by the thickness of each soil layer. Based on the average soil texture and soil organic matter content, all input soil input parameters required by the crop model were derived for the entire profile. The field capacity (SDUL - *Soil Drained Upper Limit*) and the permanent wilting point (SLL - *Soil Lower Limit*) were first estimated according to the pedotransfer function proposed by Román Dobarco et al. (2019) based on sand and clay fractions, without any consideration of soil stratification. The soil water content at saturation (*SSAT - Soil SATuration*) was estimated as the complementary fraction of the solid soil fraction, itself defined as the ratio between the soil apparent density and the mean soil particle density. Soil apparent density was estimated from soil texture according to Keller & Håkansson (2010), while the estimation of soil particle density was based on the approach proposed by Schjønning et al. (2017) and Ruehlmann (2020). Finally, the soil percolation coefficient (Kq) was estimated based on the clay content of the upper soil layer, following the approach proposed by Addiscott & Whitmore (1991). Soil AWC was estimated as :

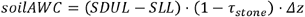

Once soil profiles were established for each STU, the frequency distribution of soil AWC was characterized within 8 × 8 km cells over the Limagne Basin (Fig. 2. A. & B.). Substantial heterogeneity for soil AWC exists between the North and the South of the Limagne plain, mostly explained by two concomitant gradients, *i*.*e*., a soil depth gradient (soil are overall shallower in the South than in the North) and a soil texture gradient (soils are more clayey in the North than in the South, at the exception of the soils along the Allier River which are sandier) (Fig. A.1). Overall, cultivated soils in Limagne exhibited a median soil AWC of 80 mm, but with considerable variability (10^th^ quantile: 40 mm; 90^th^ quantile: 118 mm). To account for the spatial heterogeneity of soils within the Limagne basin, each site was represented by the average texture of the main soils (*i*.*e*., representing at least 60% of the arable soils) of two nearby cells, which were associated with five soil depths represented by the 10^th^, 25^th^, 50^th^, 75^th^ and 90^th^ quantiles of the local zonal distribution (Fig. 2. C).

**Fig. 2.**
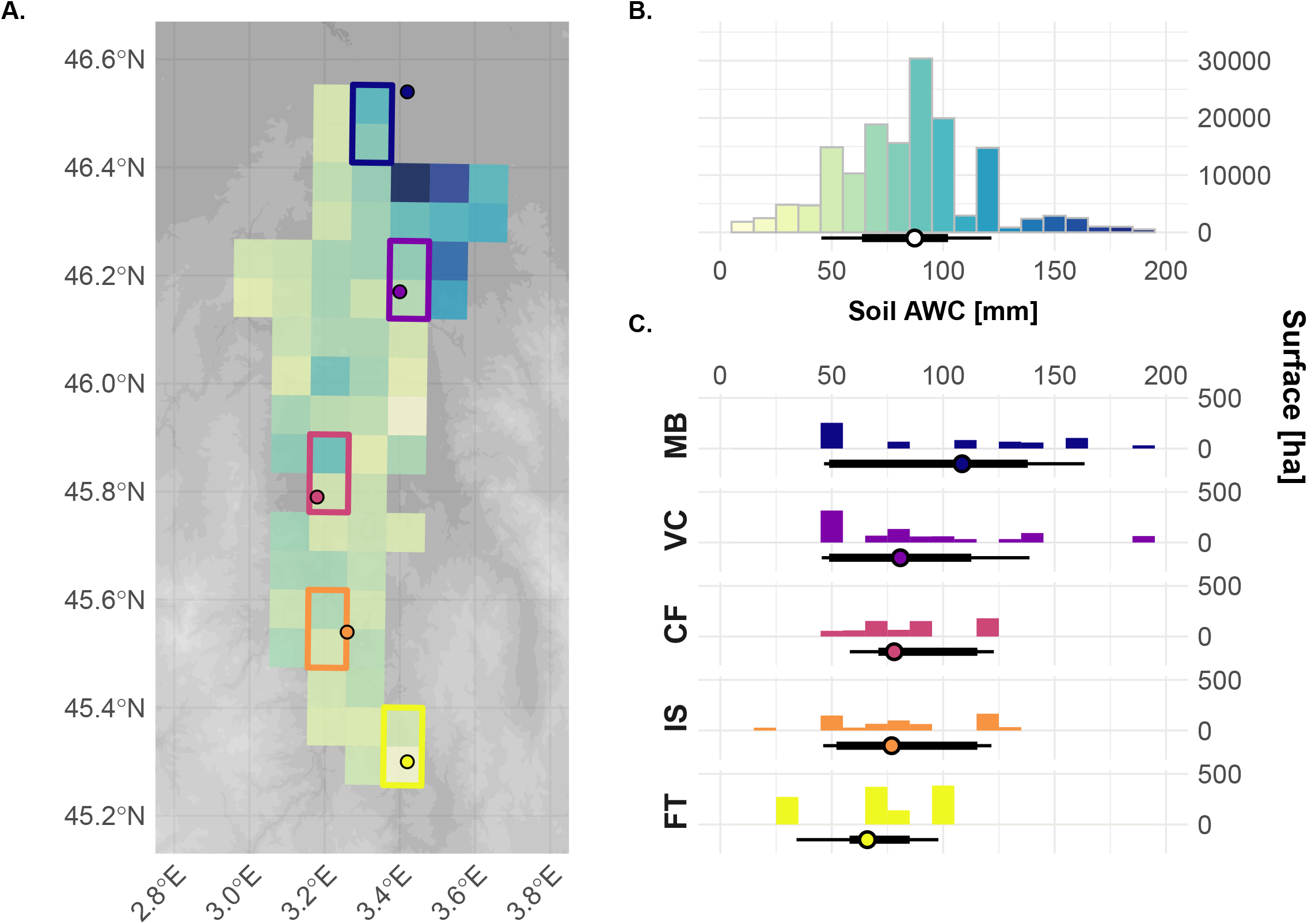
Soil available water capacity (AWC) in arable fields in the Limagne plain (A. & B.) and around each site (C.). In the gridded map (A.), each cell size is 8 x 8 km. The color scale for soil AWC is similar between the gridded map (A.) and the main histogram (B.). Under each zonal histogram (C.), the dot shows the median value, while the thick and thin lines span the interval between, respectively, the 25^th^ and 75^th^ quantiles and the 10^th^ and 90^th^ quantiles.

### Climate data

The climatic conditions prevailing over the Limagne Basin are predominantly semi-continental, largely due to the influence of local topography. The presence of mountain ranges and a dominant climatic influence of the Atlantic Ocean (on the western side) results in a pronounced Foehn effect, which is reflected in the Limagne plain being drier and milder relative to the surrounding regions. However, there are modest yet discernible climatic contrasts between the southern and northern parts of the Limagne Basin, which can be attributed to contrasting orographic sheltering. The climate of the northern part of the basin (*e*.*g*., around MB) is relatively more oceanic, exhibiting higher rainfall and smaller contrasts between temperature extremes than the southern part (*e*.*g*., around FT) according to weather observation over 1991-2020 (Fig. 1). The mean annual precipitation at MB (754 mm) differs from that at FT (612 mm) by approximately 142 mm. During the winter period, the average minimal and maximal temperatures exhibited a greater contrast in FT (-0.1°C and 10.2°C, respectively) than in MB (1.5°C and 9.4°C, respectively). This contrast is also evident during the summer months, with a greater range of average temperature extremes observed in FT (11.2°C and 26.2°C, respectively) compared to MB (13.0°C and 24.9°C, respectively).

Among the climate change emission scenarios, two shared socio-economic pathways (SSP) were considered: SSP2-4.5, representing a medium emission scenario (referred as the “*middle of the road*” scenario), and SSP5-8.5, representing a high emission scenario (referred as the “*fossil-fueled development*” scenario) (IPCC, 2023). To account for the climate change uncertainty, five global climate models (GCMs) from the CMIP6 ensemble were selected for analysis. These models were ACCESS-ESM1-5, CNRM-CM6-1, HadGEM3-GC31-LL, MPI-ESM1-2-LR and MRI-ESM2-0. Their selection leaned on a preliminary analysis conducted to identify contrasting climate trends with respect to temperature and precipitation anomaly in Western Europe. In each SSP, the temporal dynamics over the coming century were captured using three distinctive time windows, namely 2031-2050, 2051-2070 and 2071-2090. The evolution of atmospheric CO_2_ concentration in the respective scenarios and time windows is presented in Table 1.

**Table 1.** Atmospheric CO2 concentration (in ppm) according to climate change scenario and time horizon.

| Scenario | Time Horizon |  |  |  |
| --- | --- | --- | --- | --- |
|  | *1991-2020* | *2031-2050* | *2051-2070* | *2071-2090* |
| Baseline | 380.3 | - | - | - |
| SSP 2-4.5 | - | 475.4 | 538 | 584.5 |
| SSP 5-8.5 | - | 502.6 | 651 | 873.1 |

Downscaled climate data for modeling purposes were generated using a stochastic weather generator over the 5 representative sites. The downscaling procedure of the climate projections was performed using the stochastic generator LARS-WG 7.0 (Semenov, 2008) and followed the approach proposed by Semenov & Stratonovitch (2015). In essence, statistical parameters were derived for each site using observed weather datasets, which were then employed to generate 500-year weather datasets for the baseline period (1991-2020) and future time windows. Based on the statistical tests implemented in LARS-WG (Kolmogorov–Smirnov and t-tests), the weather generator showed robust performance over the baseline period. Simulated series closely matched observed monthly statistics across the five stations, with only minor and non-systematic biases (Table A.2), supporting their suitability for crop-modelling applications. We selected a weather-generator approach rather than the locally gridded downscaled Météo-France SAFRAN–DRIAS dataset (Soubeyroux et al., 2020), which has been used in national and subnational climate-change assessments (*e*.*g*. Falconnier et al., 2020; Le Roux et al., 2024). This choice was motivated by significant local biases observed over the Limagne Basin at both daily and annual scales (Quintana-Seguí et al., 2008 ; Fig. A.3).

### Model presentation

Sirius is a process-based crop model specifically designed to simulate the growth of cereal crops (Jamieson et al., 1998) and which have been validated in a wide range of environments (Ewert et al., 2002; Jamieson et al., 2000). It also has been extensively used for regional and global assessment of climate change impacts on wheat production (Asseng et al., 2015; Richter & Semenov, 2005; Semenov et al., 2014; Semenov & Stratonovitch, 2013; Senapati et al., 2021). In a nutshell, Sirius aggregates different submodels that describe water and nitrogen requirements and uptake, phenological development, and biomass accumulation and partitioning in relation with soil and climate at a daily time step. Extensive description of the model can be found in literature (Jamieson et al., 1998, 2000; Jamieson & Semenov, 2000; Lawless et al., 2005; Semenov & Stratonovitch, 2015; Senapati et al., 2019). We used Sirius version 2020 (built version 15.0.7409.24994), available at https://sites.google.com/view/sirius-wheat/. A brief description of the phenological submodel (with an emphasis on anthesis date prediction) and climate stress formalisms is provided here to ease the understanding of the proposed modeling approach.

The phenological module of Sirius describes the influence of environmental factors on vegetative and reproductive development. Respective development phases (6 distinct phases in total) are interdependent and occur concurrently. First, a pre-emergence phase is considered between sowing and leaf emergence based on fixed thermal time requirements.

Second, a leaf production phase from crop emergence to flag leaf appearance integrates the effects of vernalization requirements and photoperiod sensitivity on the emission of leaf primordia and leaf expansion. Third, the duration between flag leaf appearance and anthesis is set to three phyllochrons, while the fourth phase between anthesis and the beginning of grain filling lasts one phyllochron. The fifth phase corresponds to grain filling, whose duration is a varietal parameter defining thermal requirements. Finally, the maturation phase corresponds to 200°C degree days to allow for the grain to dry after the end of grain filling. Therefore, the anthesis date prediction is highly influenced by i) phyllochron, as a metric of the leaf emission rate in relation with accumulated thermal time, and ii) vernalization requirements and daylength sensitivity because of their impacts on the estimation of the final leaf number to be emitted. Vernalisation requirements are computed from seed moistening and daily vernalization rate increases at a constant rate (varietal parameter VAI) with daily mean soil or canopy temperature from its value at 0°C (*i*.*e*., the minimum vernalizing temperature, defined as the varietal parameter VBEE) to a maximum at 15°C (optimal vernalizing temperature). Above this temperature, the vernalization rate decreases linearly to zero until 17°C (maximum vernalizing temperature). A potential number of leaves on the main stem is determined once vernalization requirements are fully met, and an additional correction is brought to estimate the final leaf number on the mainstem depending on simulated daylength and daylength sensitivity (varietal parameter SLDL).

The effects of water limitations on wheat yield are modeled in two-ways in Sirius. First, the development rate and its incidence on plant biomass accumulation are modulated throughout the crop cycle (Jamieson et al., 1998). Second, terminal drought stress effects during reproductive development are modeled by integrating drought reduction factors on grain number and potential individual grain weight respectively (Senapati et al., 2019). With regard to the first aspect, a water stress factor (*F*_*W*_) is estimated all along the crop cycle according to the soil moisture deficit (SMD), which is defined as the difference between the actual amount of soil water in the root zone and the soil AWC in the root zone. No water stress (*F*_*W*_ = 1) is modeled until SMD reaches half of soil AWC, and increases linearly thereafter. *F*_*W*_ can decrease the rate of LAI increase during the expansion phase, accelerate the rate of leaf senescence during all phases and reduce light use efficiency in case of severe water stress. A detailed description of the related formalisms is provided by Jamieson et al. (1998). With regard to terminal drought stress, a reduction factor (R_D_) on the grain number is derived from a piecewise linear function based on a drought stress factor (DSF), which is defined as the ratio between the actual and the potential transpiration during reproductive development. The DSF is calculated as an average value over a 15-day period, starting 10 days before anthesis and ending 5 days after. The piecewise linear function R_D_ is defined by three parameters, namely DSGNR_max_ (maximum drought stress grain number reduction), DSGNS (drought stress grain number reduction saturation), and DSGNT (drought stress grain number threshold). The slope of grain reduction is then defined as S = (1 - DSGNR_max_) / (DSGNT - DSGNS). The actual number of grains per unit of ear dry matter (N_D_, in grains g^-1^) is then estimated as the product of the potential number of grains and *R*_*D*_ :

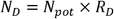

where:

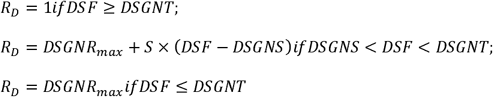

Formalisms were also incorporated to simulate the effect of thermal stress, either by accelerating the leaf senescence rate in response to high temperature or by simulating the impact of extreme temperatures (both low and high) during the reproductive phase (Stratonovitch & Semenov, 2015). Around anthesis, extreme temperatures directly impact the estimation of grain number through two reduction factors. A frost stress reduction factor (*R*_*F*_) on fertile grain number is derived from the difference between the estimated minimum canopy temperature (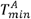 in °C) over a 6-day period (-3 to +3 days) around anthesis and a 0°C-threshold according to:

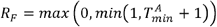

Conversely, a heat stress reduction factor (*R*_*H*_) on fertile grain number is defined from the difference between the estimated maximum canopy temperature (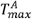 in °C) over a 10-day period before anthesis and a temperature threshold (*T*^*N*^ in °C) according to:

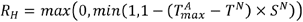

where *S*^*N*^ (°C^-1^) is the slope of the grain number reduction per unit of canopy temperature above *T*^*N*^.

Finally, the estimated number of fertile grains per unit of ear dry mass (*N*^*H*^ in grains. g^-1^) is then computed from the multiplicative product of both reduction factors

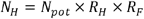

In addition, the potential seed weight can be limited by thermal heat stress when the maximum canopy temperature (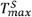 in °C) exceeds a threshold temperature (*T*^*W*^ in °C) at the beginning of grain filling (*i*.*e*., from 5 to 12 days after flowering). Then the actual weight of a single grain (W in g. grain^-1^) is estimated as follows:

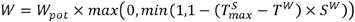

where *S*^*W*^ (°C^-1^) is the slope of the potential weight reduction per unit of canopy temperature above T^W^.

In summary, the actual yield is limited by drought and heat stresses is then calculated as:

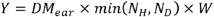

where DM_ear_ is the simulated dry matter weight of wheat ears (in g.m^-2^).

### Model instantiation and performed simulations

To isolate the effects of soil heterogeneity on wheat yield in the context of climate change, yearly crop simulations were performed with a single fixed sowing date (on the 1^st^ of November), an initial water deficit set to null (*i*.*e*., at field capacity), and non-limiting N conditions (*i*.*e*., all daily N requirements of wheat crop are fulfilled). Then, the crop simulations were performed in two steps to capture the role of breeding stress-avoidant genotypes in the context of climate change. First, a (fixed) reference genotype was used to assess the overall importance of various pedoclimatic factors on simulated yields (soil AWC, climate change uncertainty, consideration of CO_2_ increase by the model) across both baseline and future climate scenarios. In particular, the [CO_2_] effect was tested by running simulation at a constant CO_2_ level (*e*.*g*. 380 ppm taken as the mean [CO_2_] from the baseline period) or with predicted CO_2_ levels of specific SSP × time horizon combinations (see Table 1). Second, the improvement in crop performance using a stress-avoidant wheat ideotype was evaluated for each soil AWC × site under future climate change scenarios.

The fixed reference genotype was parametrized according to the cultivar Apache, which is representative of the modern cultivars currently grown in Limagne. A plant parameter file for this cultivar was obtained from previously published studies carried out within the AGMIP project (Senapati et al., 2022). Phenological parameters, related to the phyllochron (PHYLL), the photoperiod sensitivity (SLDL) or to vernalization requirements (VAI, VBEE) as well as parameters related to terminal drought sensitivity (*i*.*e*., DSGNR_max_, DSGNS, DSGNT) were recalibrated to improve predictions of both anthesis date (rRMSE = 3.77%, efficiency = 0.32) and yield (rRMSE = 14.72%, efficiency = 0.49) under current local conditions (Fig. A.4). This calibration step was performed using a self-adaptative evolutionary algorithm, as implemented in Sirius (Semenov & Terkel, 2003), on a dataset obtained from wheat variety trials conducted by INRAE in Clermont-Ferrand from 2003 to 2023 (Fig. A.4 and Table A.5).

Finally, optimal stress-avoidant wheat ideotypes were identified in each combination of site, soil depth and climate scenarios using a panel of virtual genotypes. This panel was established by optimizing three phenological parameters, namely VAI, VBEE and SLDL, to cover a range of putative anthesis dates under the current climate conditions (*i*.*e*., from DOY 100 to 160 with a regular increment of 5 days). These three parameters were chosen because, by model construction, they had no interaction effect on yield prediction for a given simulated anthesis date. Therefore, they enabled to establish a clear relationship between cultivar earliness and simulated yields (Fig. 3). In order to verify the overall validity of the approach, two distinctive yields were considered: potential yield, reflecting a theoretical maximum yield of a given cultivar under specific thermal and light conditions, and climate-limited yield, which takes into account water limitation due to precipitation scarcity or soil water depletion. The mean potential yield increased up to an optimal anthesis date, reflecting enhanced radiation interception and thermal environment, and then decreased with delayed anthesis as thermal limitations intensify (e.g., shorter grain filling) (Jamieson & Wilson, 1992). The same trend was observed for mean climate-limited yield, but with noticeable offsets: the optimal anthesis date occurred earlier than in the case of potential yield because of the occurrence of terminal stresses (Flohr et al., 2017; Zheng et al., 2012). To prevent the introduction of irrelevant parameter values, the range of parameters explored by the optimization algorithm was constrained according to published studies (Table A.5). A total of 90 virtual wheat genotypes were established and used in every environment to identify the “optimal” genotype, *i*.*e*., the one that maximizes mean yield in a given situation (site × soil AWC × time horizon × GCM × SSP).

**Fig. 3.**
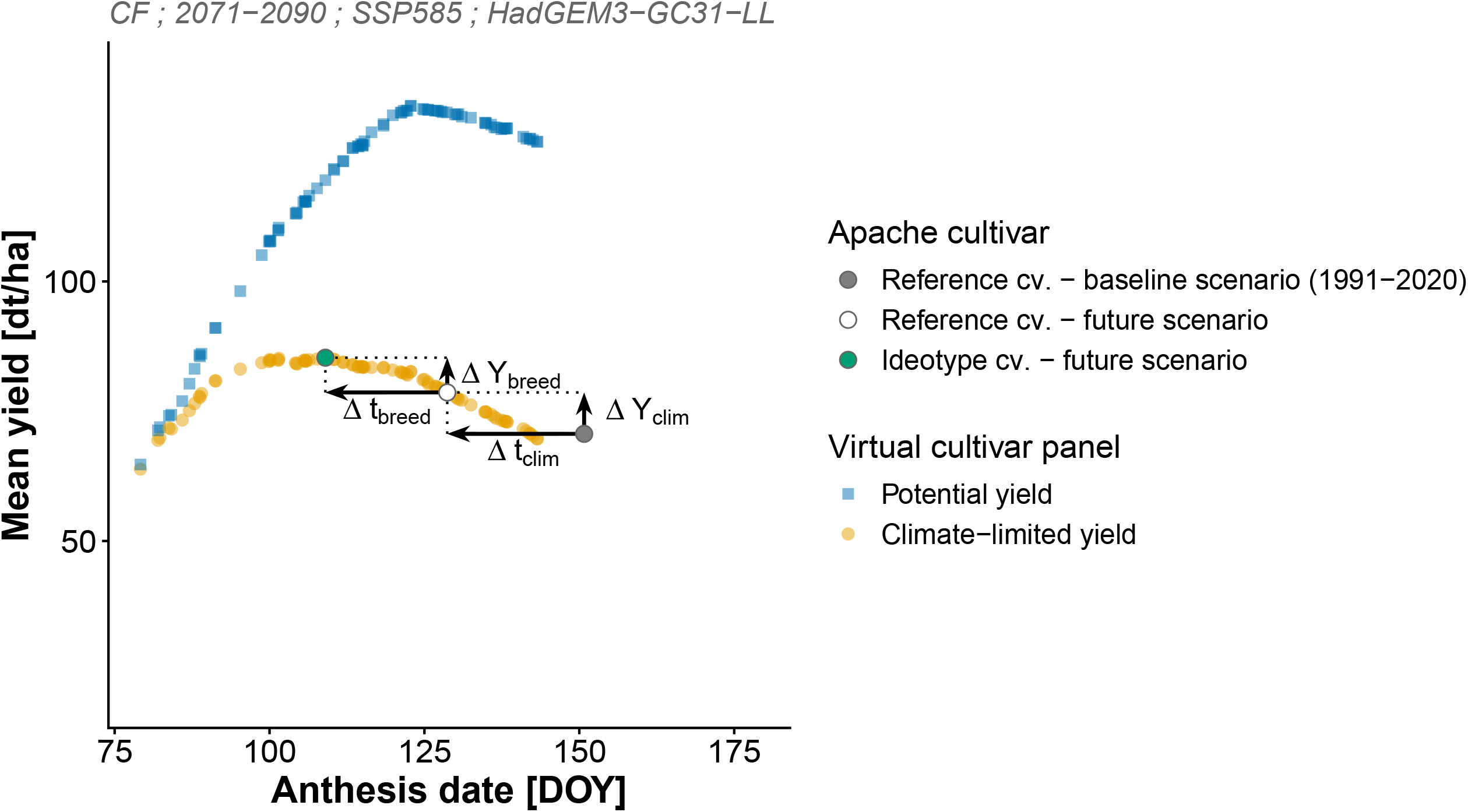
Illustrative example of projected climate impacts on the mean yield of a virtual panel of wheat genotypes over a wide range of anthesis dates. In each situation (RCP x GCM x time horizon x location x soil depth), the respective effects of i) climate change (X_clim_) and ii) an optimal stress-avoiding breeding strategy (X_breed_) are assessed according to the changes in mean anthesis date (Δt) and mean yield (ΔY) of two Apache cultivars: a reference one with current phenological parameters (in baseline and future conditions) and an optimal one, with an adjusted set of phenological parameters.

### Data analysis

To gather insights on the climate modifications induced by climate change, seasonal anomalies for both mean temperature and precipitation were computed. The period 1991-2020 was used as a baseline period, and seasonal anomalies were established for all GCMs, SSPs, and time horizons. To illustrate the temporal evolution of these anomalies, trajectories were represented graphically: arrows connect successive time horizons, with nodes marking the anomaly values at each horizon (e.g., 2031–2050, 2051–2070, 2071– 2090). The position of the nodes highlights the anomalies specific to each horizon, while the direction and length of the arrows depict both the trend and the magnitude of projected changes over time (see Fig. 4 for absolute anomalies and Fig. A.6 for relative anomalies).

**Fig. 4.**
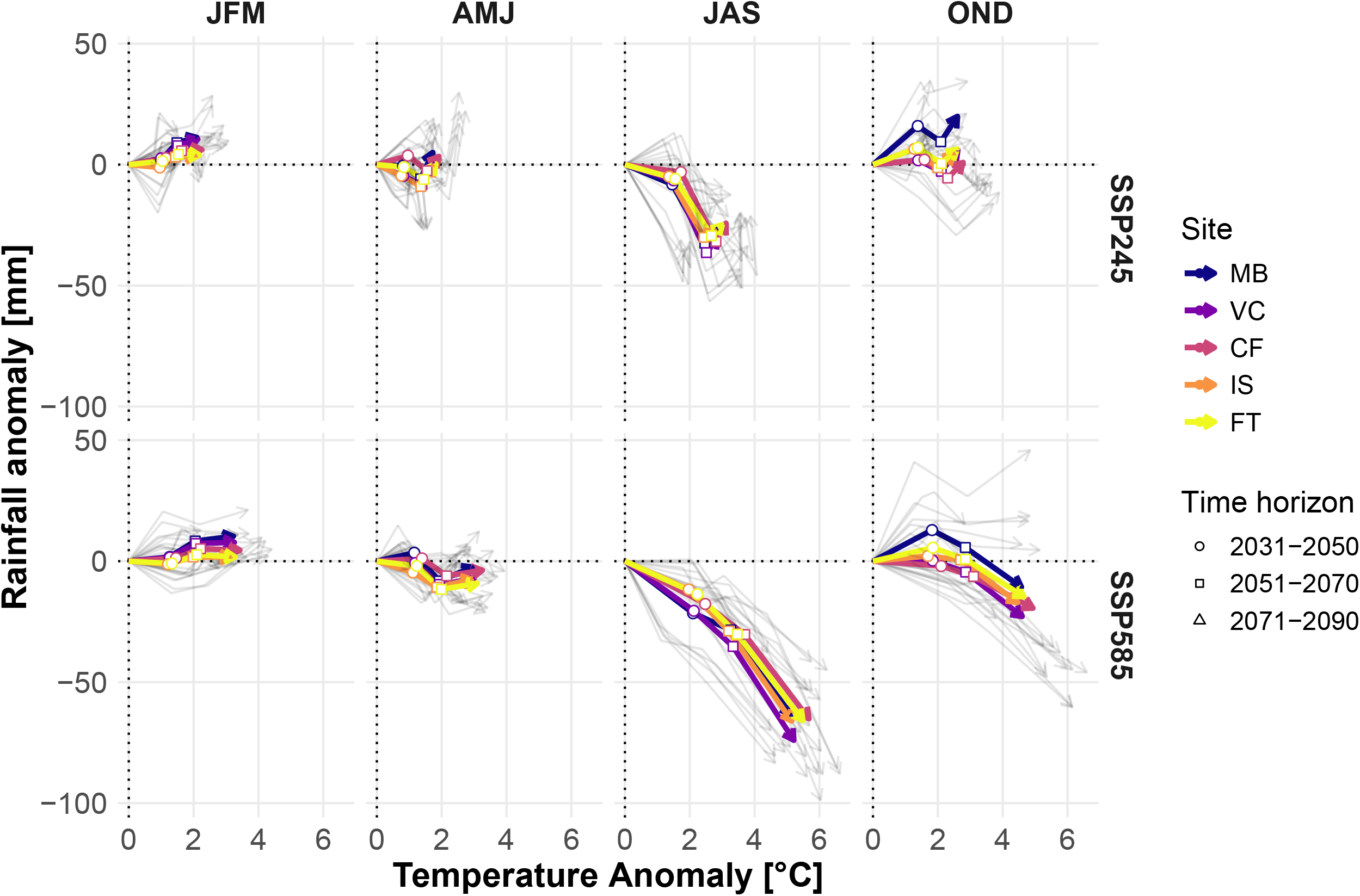
Seasonal trajectories of absolute temperature and precipitation anomalies for each site. Anomalies were calculated relative to the baseline period (1991-2020). Each arrow node corresponds to the coordinates of the anomalies at a given time horizon. Thick colored arrows show the median trajectories for each site while thin gray arrows show the dispersion of trajectories for each site-GCM combination. Trajectories of relative anomalies are provided in Appendix S6.

The influence of pedoclimatic factors on simulated wheat yields was evaluated across time by applying an analysis of covariance (ANCOVA). Explanatory factors included site, soil AWC, SSP, GCM, and crop response to atmospheric CO□ ([CO□]). By construction of the statistical model, residuals represented mainly the remaining inter-annual climate variability. For the reference period (1991–2020), only site and soil AWC were considered, whereas future periods additionally included SSP, GCM, and CO□. Candidate additive models, with and without selected two-way interactions, were compared using Akaike’s Information Criterion (AIC), and the model with the lowest AIC was retained. To ensure comparability across future horizons, a single common model was selected based on the best overall fit (lowest sum of AIC across horizons) and applied consistently. Then, variance decomposition was performed on the final ANCOVA models to quantify the contribution of each factor (fixed and residual) to total yield variability across all periods.

To quantify the impacts of climate change, the performances of the reference genotype were quantified in terms of absolute change of both yield and anthesis date. Climate change effects were quantified in all sites × soil AWC × GCM × SSP combinations according to the mean phenological shift (Δt_clim_, in days) and mean yield modification (ΔY_clim_, in dt. ha^-1^). Both metrics were computed relatively to a baseline scenario (1991-2020) (Fig. 6), and calculated as:

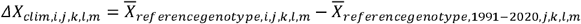

where X is the mean variable of interest (Y or t), *i* is a future time horizon, *j* a given site, *k* a soil AWC quantile, and *l* and *m* are respectively the indices of specified GCMs and SSPs.

The effects of an optimized breeding strategy were evaluated similarly, in terms of phenological shift (Δt_breed_, in degree. days^-1^) and yield gain (ΔY_breed_, in dt. ha^-1^) relative to the reference genotype (Fig. 7) in a given environment. It was calculated as:

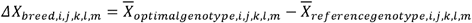

where X is the mean variable of interest (Y or t), *i* is a future time horizon, *j* a given site, *k* a soil depth quantile, and *l* and *m* are respectively the indices of specified GCMs and SSPs.

To assess the statistical significance of simulated yield changes, paired Wilcoxon tests were applied to yearly yield series, both for climate change effects (ΔY_clim_) and for breeding effects (ΔY_breed_), across all site × soil × GCM × SSP combinations. This non-parametric test evaluated whether yield distributions differed significantly between contrasted scenarios (baseline vs. future, or reference vs. optimized genotype), with significance considered at *p* ≤ 0.01.

Finally, the exposition of the fixed reference genotype and all stress-avoidant ideotypes to frost risk at 1 cm ear stage was evaluated from a probabilistic perspective, since such climate risk is not considered by Sirius though its increasing importance for designing future ideotype is assessed in existing literature (Bogard et al., 2021; Zheng et al., 2015). The frost exposure frequency was defined as the occurrence probability of detrimental frost temperature (daily minimum temperature ≤ -5°C) after the initiation of stem elongation (or 1-cm ear stage, GS30) (Bogard et al., 2021). Since Sirius does not provide any information about the initiation of stem elongation, this stage was estimated considering a fixed period of 500°C.day prior the anthesis date based on available literature (Roychowdhury et al., 2023; Whitechurch et al., 2007). The frost exposure frequency was estimated for the fixed reference genotype and all stress-avoidant ideotypes in each site × soil depth × time horizon × GCM × SSP.

All analyses were conducted in R (R Core Team, 2020). Data wrangling, statistical modeling, and figure generation relied on a combination of community-contributed packages available through CRAN.

## Results

### Hotter years and drier summers are forecasted in Limagne

The seasonal temperature anomalies are expected to increase during the upcoming century, with the magnitude of these changes being dependent on the SSP (Fig. 4 and A.6). By 2071-2090, the mean annual temperature anomaly is projected to reach +2.4 to +4.0 °C according to SSP2-4.5 and SSP5-8.5, respectively. While the temperature anomaly will increase with time during all seasons, the warming trend is most pronounced during the summer (+2.9 to +5.3 °C) and fall (+2.6 °C to +4.7 °C) periods.

Precipitation anomalies are projected to differ strongly between seasons, with summer precipitation potentially declining significantly over time (Fig. 4 and A.6). By 2071-2090, all GCMs predict a decrease in summer precipitation, with an average reduction of -13.4 % (-27 mm) and -33.7 % (-67 mm) according to SSP2-4.5 and SSP5-8.5, respectively. Oppositely no clear trend appears when other seasons are considered, since precipitation modifications are low in magnitude but also inconsistent between GCMs. For example, winter precipitation is anticipated to increase gradually, but reaching only + 6.5 % (+ 7 mm) and + 4.6 % (+ 5 mm) according to respective SSP scenarios. Trajectories of precipitation anomalies over the spring and fall seasons indicate minimal change over the course of the upcoming century, with noticeable discrepancies among the GCMs. At an annual scale, the overall climate tends to become slightly drier by the end of the century (on average -10 to -85 mm depending on SSP).

Finally, the projected climate trajectories exhibited at the five investigated sites are similar. An analysis of climate trajectories according to temperature anomalies revealed no significant differences between the northern and southern parts of the Limagne basin. Slight differences were observed for precipitation anomalies during the fall and winter seasons, but given the observed variability of GCMs, the degree of uncertainty remains considerable.

### Soil heterogeneity at regional level is the major source of variability of simulated yield

The importance of local soil heterogeneity on simulated yield is reflected by a larger range in northern sites (*i*.*e*., MB and VC) were larger panels of soil AWC exists in comparison to southern sites (*i*.*e*., IS and FT) (Fig. 5.A). For instance, mean simulated yield ranges between 53.1 and 92.0 dt. ha^-1^ in MB (soil AWC: 44-164 mm) while it only ranges between 43.4 and 60.5 dt. ha^-1^ in FT (soil AWC: 43-99 mm). At a given site, higher soil AWC tends to reduce interannual yield variability. In MB, standard deviation of simulated yield decreases from 17.9 dt. ha^-1^ (CV = 33.7%; soil AWC = 44 mm) to 11.9 dt. ha^-1^ (CV = 11.9 %; soil AWC = 164 mm). Nevertheless, the reduction of the interannual yield variability along with soil AWC is not systematic. In FT, where the range of soil AWC is narrower, the yield variability remained relatively constant between shallow (σ = 16.9 dt. ha^-1^; CV = 39.0 %; soil AWC = 43 mm) and deep soils (σ = 20.6 dt. ha^-1^; CV = 35.0 %; soil AWC = 99 mm), suggesting either local climate or threshold effects related to soil AWC that influence interannual yield variability.

**Fig. 5.**
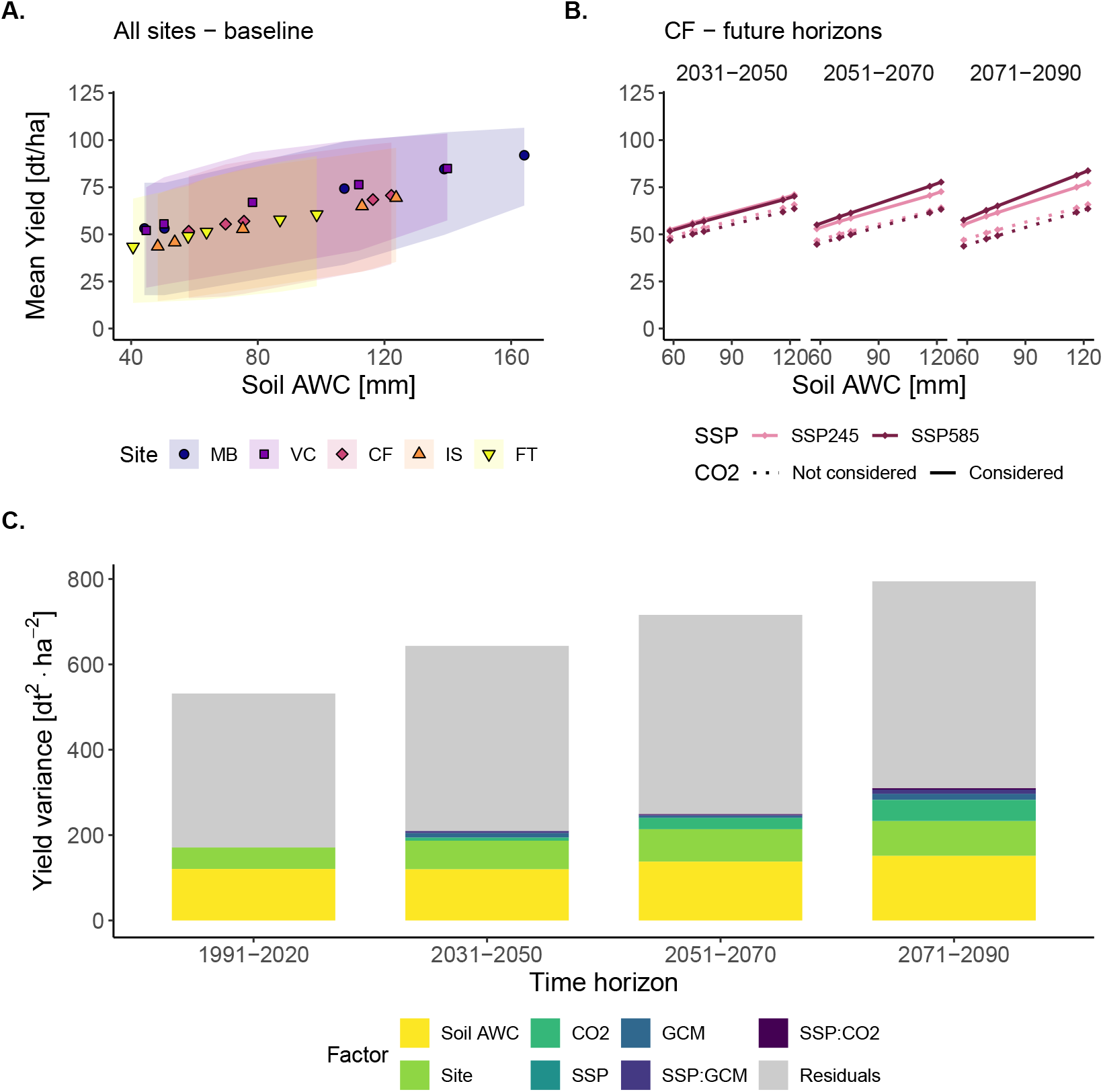
Contribution of soil AWC to simulated wheat yields, for the reference genotype across all sites (A.), at Clermont Ferrand (CF) in future time horizons (B.) and the overall variance decomposition according to ANCOVA for current and future time horizons (C.). The relationship between mean wheat yield and soil AWC is depicted for all sites under the baseline scenario (1991-2020) (A). The envelope curves show the range of variation between the 10^th^ and 90^th^ quantile of the simulated yields for each site.

According to SIRIUS, the contribution of [CO_2_] to yield increase was significant, but the absolute contribution differed among soils due to contrasting yield potential (Fig. 5.B. & 5.C.). Separate consideration of climate sequences and CO_2_ concentrations revealed that simulated yield stagnated or decreased slightly under constant [CO_2_] (*i*.*e*., 380 ppm). Using CF as an example (with a mean simulated yield of 60.6 dt.ha^-1^ in 1991-2020), the simulated yields under constant [CO_2_] amounted to an average of 56.7, 54.9 and 56.0 dt.ha^-1^ in 2031-2050, 2051-2070 and 2071-2090, respectively, with the climate sequences of the SSP2-4.5 scenario. With the climate sequences of the SSP5-8.5 scenario, the respective yields were 54.8, 53.5 and 53.2 dt.ha^-1^. Conversely, when considering the projected increase of [CO_2_] in the SSP scenarios, the average simulated yields increased substantially and progressively. For instance, under the SSP5-8.5 scenario and its forecasted [CO_2_] evolution, mean simulated yields reached 60.5, 65.8 and 70.1 dt.ha^-1^ in 2031-2050, 2051-2070 and 2071-2090, respectively. More broadly, these trends were simulated across all sites (Fig. A.7), suggesting that the predictions of future yield improvements were intrinsically related to the formalisms used to capture the effects of [CO_2_] on plant biomass accumulation. Finally, simulated absolute yields improved significantly in deep soils, whereas improvements related to [CO_2_] were more limited in shallower soils. Taking the predictions at CF under the SSP5-8.5 scenario as an example, the increase related to [CO_2_] in deep soils was +6.50, +14.30 and +19.9 dt.ha^-1^ in 2031-2050, 2051-2070 and 2071-2090, respectively. In contrast, the increase in shallow soils was +5.16, +11.00 and +15 dt.ha^-1^ in the same periods. No differences were observed between soils on a relative basis, suggesting that CO_2_ fertilization exacerbates the yield potential independently of soil depth. These results were also observed more generally across all sites (Fig. A.8).

Variance decomposition revealed that, at the scale of the Limagne territory, soil AWC is the most important pedoclimatic factor among selected sites, while site effects, mainly driven by climate contrasts at the territory scale, were a secondary explanatory factor (Fig. 5.C). Under current climate conditions, soil AWC explained 22.7% of the variance while site effects amounted only for 9.5%. In future climate conditions, the overall yield variability increased due to climate change uncertainty (SSP, GCM), and increased [CO_2_]. However, the respective proportional influences of soil AWC (18.6%, 19.2% and 19.0% in 2031-2050, 2051-2070 and 2071-2090 respectively) and sites (10.4%, 10.6% and 10.3% respectively) remained stable by the end of century. On the contrary, the proposed decomposition showed an increased influence of the [CO_2_] (1.1%, 3.7% and 6.2% in 2031-2050, 2051-2070 and 2071-2090 respectively), although it remained minor relatively to the role of local soil or site-related climate contrast. The contribution of climate models (GCMs, SSPs and their interaction components) was very limited at the scale of the study area. Last but not least, residual variability remained the primary source of yield variability (67.8%, 67.4%, 65.1% and 61.0% in 1991-2020, 2031-2050, 2051-2070 and 2071-2090 respectively), highlighting the importance of interannual climate variations.

### Hastened phenology and soil dependence of future yields

The anthesis date of the fixed reference genotype will gradually become earlier over the next century, with the magnitude of this shift depending on the SSP considered (Fig. 6). In SSP2-4.5, the mean anthesis date is projected to be 8.6 to 16.6 days earlier between 2031-2050 and 2071-2090 compared to the baseline. In SSP5-8.5, it is projected to be 11.2 and 21.4 days earlier, respectively.

**Fig. 6.**
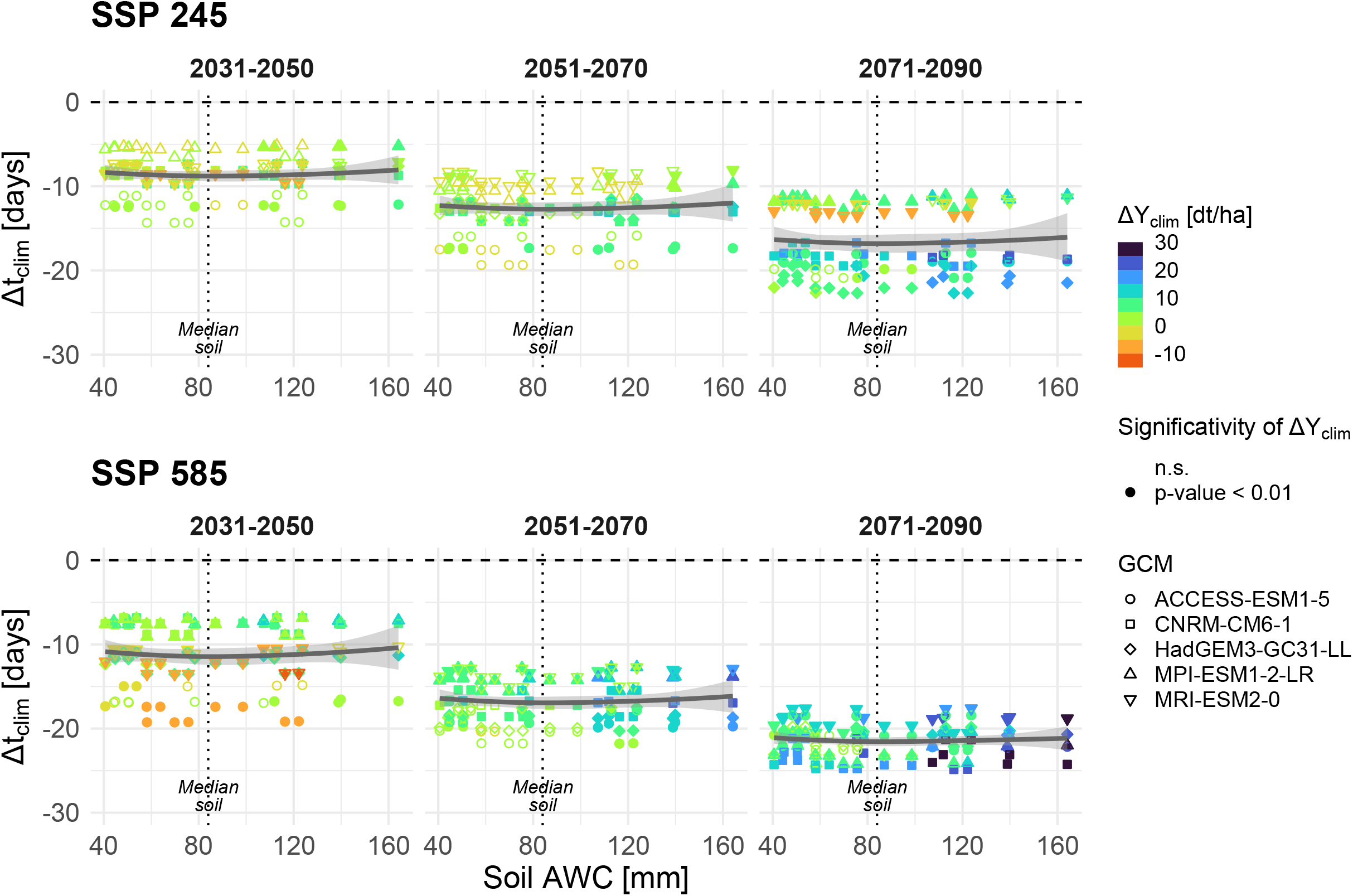
Decomposition of the impacts of climate change on a reference genotype (*cv*. Apache) in terms of anthesis date (Y-axis) and crop yield (color) in relation with soil available water capacity (X-axis). A loess smoothing depicts the overall trend of phenology shift along the soil AWC. Statistical significance of the mean difference in terms of crop yield is represented by filled and hollow signs.

As the century progresses, the simulations indicate an overall improvement in wheat yields, regardless of soil AWC. By 2031-2050, the yield increase is projected to reach 1.16 and 2.13 dt. ha^-1^ in SSP2-4.5 and SSP5-8.5 respectively, while it further increases to 7.55 and 12.3 dt. ha^-1^ by 2071-2090. Over the entire century, the simulations suggest that the yield increase observed for the same reference genotype is more pronounced in deep soils than in shallow soils. Using the mean soil AWC of Limagne (80 mm) as a discriminative, yet subjective, threshold between deep and shallow soils, the mean yield increase in deep soils is about twice that of shallow soils in each time horizon and SSP.

### Influence of local soil heterogeneity on the earliness of stress-avoiding ideotypes

The thermal time required to reach the mean anthesis dates of optimal stress-avoiding ideotypes depends on soil AWC according to the model prediction (Fig. 7). Averaging all time horizons, GCMs and SSPs, the model suggests that the mean Δt_breed_ is about -102 °C.days (σ = ± 96.4 °C.days) in shallow soils and -214 °C.day (σ = ± 68.3 °C.days) in deep soils. In addition, the model predictions suggest that Δt_breed_ along the soil AWC gradient is relatively similar across all time horizons within a given SSP scenario. For the SSP2-4.5 scenario, the required thermal time for wheat ideotypes in shallow soils is on average -131, - 132 and -77 °C.days by 2031-2050, 2051-2070 and 2071-2090 respectively, while it is -249, -217 and -173 °C.days in deep soils. Similarly, the average required thermal time in shallow soils is -66, -115 and -90 °C.day in the SSP5-8.5, and it amounts to -228, -212 and -203 °C.days in deep soils.

**Fig. 7.**
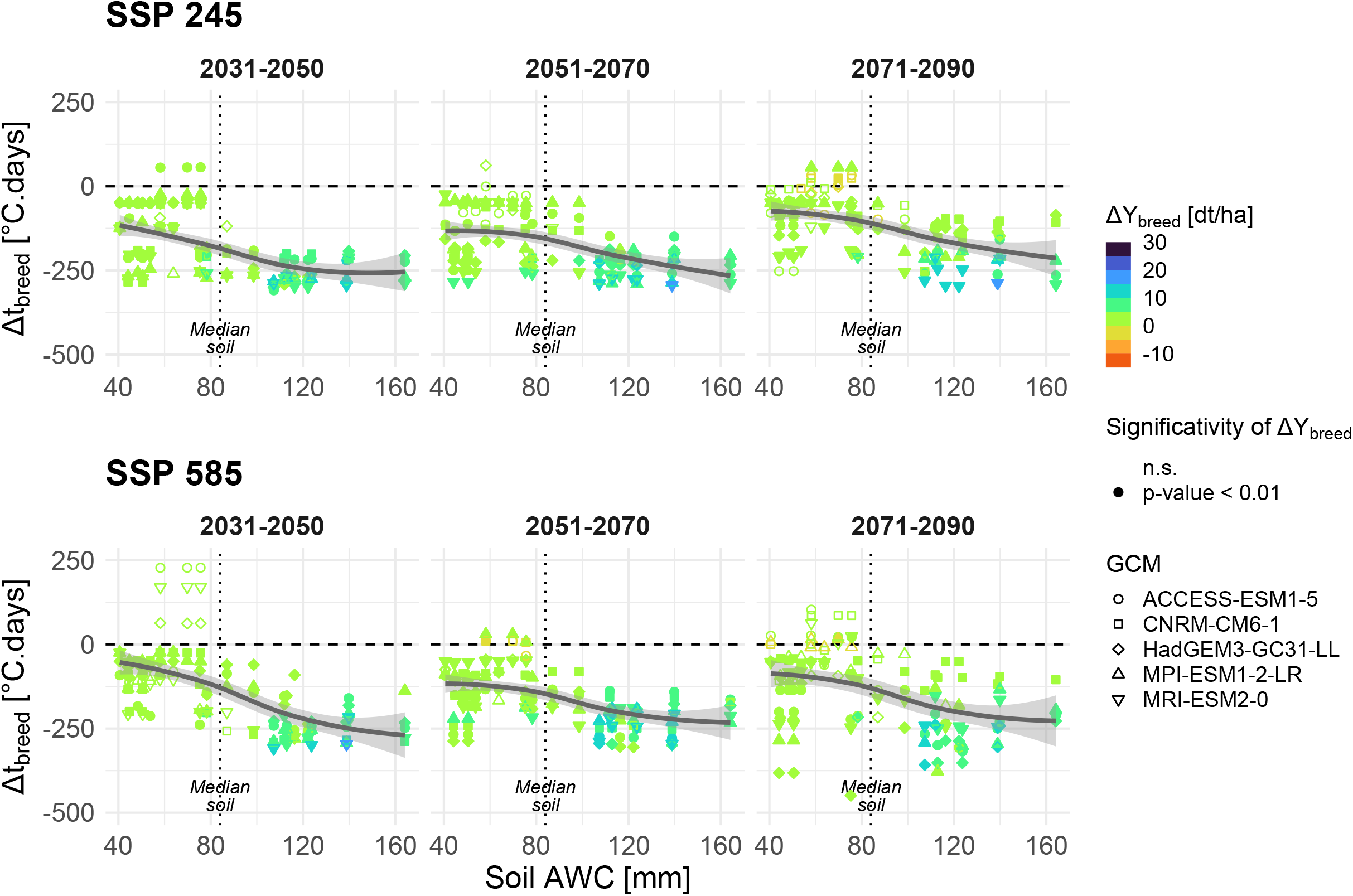
Decomposition of the theoretical breeding gain in terms of anthesis date (Y-axis) and yield (color) in relation with soil AWC (X-axis). The theoretical breeding gain is established by comparing a phenologically-optimized ideotype and a reference genotype (Apache *cv*.). A loess smoothing was used to represent the thermal time required to breed optimal stress-avoiding genotypes along the soil AWC. Statistical significance of the mean difference in terms of crop yield is depicted by filled and hollow signs.

Similar to the predictions obtained with the reference genotype, the magnitude of the associated yield gain with optimal stress-avoiding ideotypes is higher in deeper soils than in shallower soils. Averaging all time horizons, GCMs and SSPs, ΔY_breed_ amounts on average +6.33 dt. ha^-1^ in deep soils (σ = ± 3.52 dt. ha^-1^) and +1.71 dt. ha^-1^ (σ = ± 1.39 dt. ha^-1^) in shallow soils. Similar to the predictions of Δt_breed_, mean ΔY_breed_ are similar across all time horizons within a given SSP scenario. For the SSP2-4.5 scenario, ΔY_breed_ in deep soils is on average +7.40, +7.24, +5.03 dt.ha^-1^ by 2031-2050, 2051-2070 and 2071-2090 respectively, while it reaches +2.02, +2.16, +1.24 dt.ha^-1^ in shallow soils. In the SSP5-8.5 scenario, mean ΔY_breed_ is overall lower and amounts in deep soils to +5.98, +6.99, +5.33 dt. ha^-1^ by 2031-2050, 2051-2070 and 2071-2090 respectively, while it accounts in shallow soils for +1.33, +2.08, +1.44 dt. ha^-1^.

### Early-spring frost exposure and intra-regional disparities

The probability of frost occurrence at the 1-cm ear stage remained overall stable for the reference genotype in both SSP scenarios but revealed substantial regional disparities (Fig. 8-A). In the northern sites (MB, VC and CF) with a milder climate, the probability of early frost events remained low (P_frost_ ≤ 0.05, *i*.*e*., less than 1 year out of 20) and stable over the century. On the opposite, the southern sites (IS and FT) exhibited a higher probability of frost event at 1 cm ear stage (P_frost_ = 0.096 and 0.076 respectively in 1991-2020). In these sites, an increase in frost probability was predicted toward the end of the century, particularly under SSP245 (P_frost_ = 0.175 and 0.116 respectively in 2071-2090), whereas stronger warming under SSP585 led to a limited increase at IS (P_frost_ = 0.140 in 2071-2090) or a small decrease at FT (P_frost_ = 0.048).

**Fig. 8.**
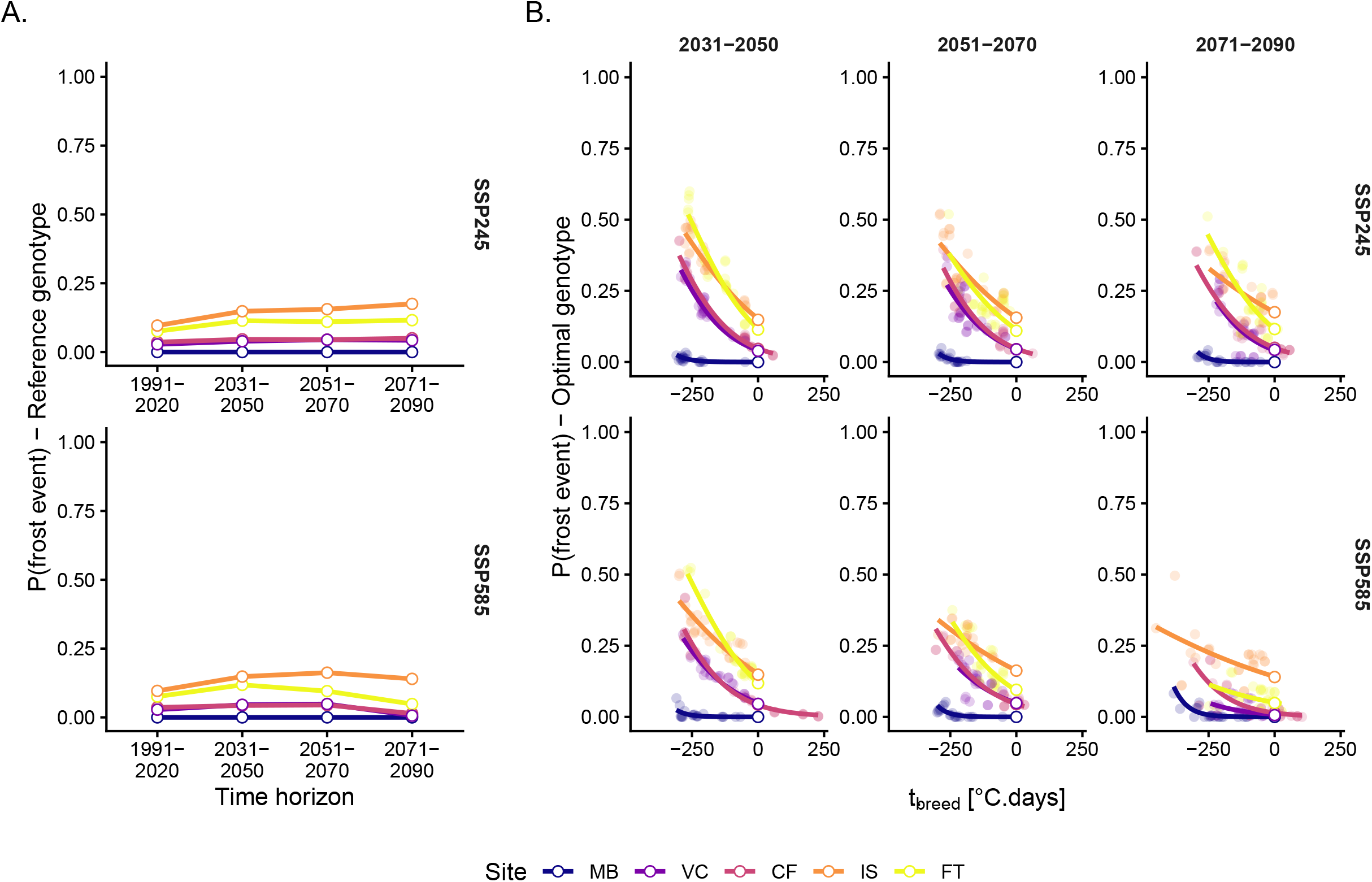
Temporal evolution of frost exposure risk after 1-cm ear stage for reference genotypes (A.) and for stress-avoidant ideotypes according to their theoretical breeding gain (B.). Optimal genotypes in various environments are depicted with filled circles while reference genotypes are represented with hollowed circles.

When considering breeding-induced shifts in phenology (Fig. 8-B), frost risk followed inverse exponential pattern at respective sites. Relative to the reference genotype (Δt_breed_ = 0 °C.d), advancing flowering increased frost probability sharply: around Δt_breed_ = 100 °C.d, the mean frost risk probability increased between 67% and 103% in SSP2-4.5 (depending on the considered period) and between 67% and 97% in the SSP5-8.5. Conversely, the average frost risk increased between 295% and 401% in the SSP2-4.5 (depending on the considered period) and between 264% and 344% in the SSP5-8.5 when flowering occurred earlier at Δt_breed_ = -250 °C.d. The increase rate of frost exposure to Δt_breed_ appeared to be strongly site dependent, with steeper increase in the frost-prone sites (IS and FT) than in the milder locations (VC and CF). MB showed a distinctive pattern with very limited risk increase even for the most hastened genotypes (P_frost_ < 0.010 for Δt_breed_ < -250°C in all SSP × GCM × time horizon conditions). Overall, these results suggest that frost risk responds asymmetrically to phenological shifts, with stronger sensitivity to flowering advancement in frost-prone sites.

## Discussion

### Intra-regional soil heterogeneity as a dominant driver of future yield variability

As expected, our simulations show that intra-regional soil heterogeneity is a dominant driver of wheat yield variability at the regional scale. Variance decomposition performed on our simulations showed that, soil available water capacity (AWC) explained up to 23% of simulated yield variability, which is twice the contribution of site-related climate contrasts (10%, Fig. 5.C.) within the study area. If inter-annual variability is let aside, soil AWC is the main cause of variance in our simulations. In addition, the influence of soil AWC persists over time horizons, thus confirming that this factor remains crucial in yield determination in spite of the growing effects of climate change on inter-annual variability.

Beyond our case study, similar patterns have been reported in other European contexts through modeling studies, suggesting that this dominance of soil heterogeneity is not site-specific. In a region from southern Norway, Persson & Kværnø (2017) found that taking into account spatial soil heterogeneity by using detailed soil information did not alter mean wheat yields but strongly increased the range of simulated yields across spatial units. Similarly, in northern Germany, Hoffmann et al. (2016) showed that aggregating soil and climate inputs substantially reduced simulated yield variability, thereby masking fine-scale spatial heterogeneity and underestimating local yield contrasts. Together, these results highlights that an explicit representation of the variability of soil properties is essential in climate impact assessments, in order to avoid any underestimation of local climate-driven water-limitation effects that would, in return lead to overestimate putative adaptive potential.

### Soil AWC shapes wheat yield trajectories under climate change

An explicit accounting of soil AWC variability is also required when assessing the yield impact of wheat phenology modifications driven by climate change.

As widely observed in other studies (Porter & Gawith, 1999; Rogger et al., 2021; Semenov, 2009; Zheng et al., 2016), the first consequence of the gradual warming induced by climate change is the overall acceleration of wheat phenology. In the present study, when considering a reference genotype, the mean advancement of anthesis ranged from 8.6 to 21.4 days compared to the baseline, depending on the time horizon considered and the SSP scenario. The shorter crop cycle that results naturally implies a reduction in crop radiation interception, but this negative effect on biomass accumulation is counterbalanced by the gradual rise in [CO_2_]. Additionally, the acceleration of wheat phenology induced by climate change tends to limit wheat’s exposure to terminal stresses, thereby limiting their negative impact on yield. Therefore, the effects of climate-driven phenology acceleration on yield are governed by processes that respond differently to soil AWC, again highlighting the crucial importance of soil input data.

In the present study, this interaction is the main cause of a marked divergence in yield trajectories between shallow and deep soils under future climates. In low-AWC soils (40–80 mm), mean yields of a reference genotype are projected to stagnate or decline, whereas in high-AWC soils (80–160 mm), yields tend to increase (Fig. 6). Intuitively, one may anticipate that enhanced wheat earliness driven by climate change may benefit more to crops on shallow soils that are more prone to frequent and intense drought stress caused by the lack of water reserve in those soils. This is not the case. This appears to be caused by the strong buffering capacity of deep soils that allow to potentialize the projected fertilization effect of elevated [CO_2_] over the whole crop cycle. Indeed, the stimulation of photosynthesis by elevated [CO_2_] enhances potential carbon assimilation, but its translation into yield gain depends on the plant’s ability to maintain growth and grain filling. When soil moisture is adequate, the CO_2_-induced improvement in water-use efficiency sustains canopy development and biomass accumulation, amplifying yield gains (Fitzgerald et al., 2016; Kimball, 2016; O’Leary et al., 2015).

By contrast, on shallow soils, increasingly frequent water deficit, even of mild intensity, limit the expression of the CO_2_ fertilization effect on biomass accumulation and do not allow to translate the fastening of the crop cycle into sensible yield gain in spite of the avoidance of severe terminal stresses. Therefore, accounting for local soil heterogeneity is a key requirement for the definition of regional ideotypes.

### Implications of soil heterogeneity for regional climate risk assessments

A key message of our study is that ignoring soil heterogeneity may distort climate risk assessments at larger scales. At national or regional scales, climate risk assessments of crop production often rely on aggregated indicators, in which water deficit is inferred from precipitation–evapotranspiration balances (Beauvais et al., 2025; Le Roux et al., 2024). While informative, these indicators ignore soil buffering capacity and may misrepresent actual stress exposure: risks may be underestimated in shallow soils while overestimated in deep soils.

Our results further demonstrate that this bias may occur even within relatively homogeneous climatic regions, as illustrated in the Limagne basin, where soil AWC dominates over climate contrasts. Furthermore, soils with limited AWC are widespread, as nearly half of local soils exhibit AWC ≤ 80 mm, and this constraint is not restricted to a few marginal areas but affects large parts of the basin beyond the deep vertisols around Clermont-Ferrand. Such intra-regional soil contrasts are unlikely to be confined to Limagne and are expected to occur across many European wheat-growing regions.

Investigating such intra-regional soil-driven biases at broader scales requires spatially explicit soil information. In this respect, France provides an instructive context, as harmonized soil and climate databases are increasingly available at national and sub-national scales (Debaeke et al., 2017; Lémond et al., 2011; Voltz et al., 2020). However, key uncertainties remain regarding soil thickness which largely determines rooting depth and effective AWC (Arrouays et al., 2020; Cousin et al., 2022).

These uncertainties have direct methodological implications for climate change impact assessments. Continental-scale assessments often rely on sparse site networks assuming a single soil profile per location, thereby limiting the transferability of their conclusions to the regional scale. As illustrated by the Clermont-Ferrand case, representing a region using a single, average soil profile tends to bias simulations toward deep, favorable soil conditions, ultimately masking contrasted yield responses across soil profiles.

### Ideotyping stress-avoidant genotypes depends on local soil available water content

Therefore, we argue that future ideotyping strategies cannot be generalized at the (sub-) regional scale without explicit consideration of soil profile variability. Stress-avoidant cultivars optimized for deep soils may indeed benefit from greater water buffering and potentially from CO_2_ fertilization, whereas the same ideotypes may perform poorly in shallow soils, where limited gain are projected (Fig. 7).

Another key finding of our study is that the thermal breeding requirements and the expected yield gains of stress-avoidant ideotypes are dependent on soil AWC. In deep soils, advancing anthesis by about 150–200 °C days consistently improved simulated yields (Fig. 7). This confirms that early-flowering genotypes can benefit from escaping terminal drought and heat when sufficient soil moisture remains available (Flohr et al., 2017).

Under these conditions, greater soil storage sustain biomass accumulation and grain filling despite a shorter cycle. In contrast, in shallow soils (AWC < 80 mm), the proposed ideotyping procedure suggests lower thermal breeding requirements to optimize mean yield, and the expected yield gain appears more limited. In other words, the stress-avoidant breeding strategy has limited capacity to adapt to future climate conditions in shallow soils.

These results indicate that a single phenological target within the same region cannot serve as a universal ideotype. In regions such as Limagne, where drought is projected to remain a dominant climatic constraint (Beauvais et al., 2025; Le Roux et al., 2024; Semenov & Shewry, 2011), defining optimal anthesis timing in relation to local soil water availability is likely more appropriate. Building on previous evidence that accounting for cultivar shifts is essential to realistically assess climate change impacts on wheat phenology (Rezaei et al., 2018), our findings indicate that adaptive breeding priorities should also be differentiated along pedological gradients. In deep soils, phenological stress avoidance remains an effective strategy, potentially complemented by traits improving root exploration or canopy resilience under moderate stress (Christopher et al., 2016; Wasson et al., 2012). In shallow soils, opportunities for avoidance appear more limited; and breeding efforts should instead prioritize tolerance traits that enhance grain-filling stability under both drought and heat (Li et al., 2022; Senapati et al., 2026; Tricker et al., 2018). Given the inherently constrained potential yield of these environments, breeding for yield stability rather than maximum productivity may also be an effective strategy.

### Reconciling terminal stress avoidance with frost tolerance

Finally, one should not forget that shifting phenology toward earlier anthesis may also advance the exposure of reproductive stages to late frost events. Integrating frost tolerance into ideotyping frameworks will therefore be essential to balance terminal-stress avoidance with early-season cold risks under future climates.

Despite ongoing climate warming, damaging frosts in late winter and early spring are expected to remain a key constraint for phenological adaptation strategies across Europe. Climate change modifies the alignment between crop phenology and frost occurrence, meaning that earlier genotypes designed to avoid terminal heat and drought may face higher frost exposure (Zheng et al., 2012, 2015; Bogard et al., 2021).

At the regional scale, ideotyping strategies aimed at escaping terminal stress cannot be generalized without accounting for spatial variability in frost exposure. Frost risk remains heterogeneous, shaped by elevation, topography, and local climatic conditions. The spatial drivers of frost risk differ from those governing soil heterogeneity, implying that both constraints must be jointly considered rather than addressed independently. Consequently, while the development of early-flowering ideotypes is a relevant option to avoid terminal stresses, it must be combined with sufficient frost tolerance to ensure crop stability in frost-prone areas.

In Limagne, both current and future climates exhibit marked spatial contrasts in frost exposure, with persistent frost in specific areas of the plain (especially in the South which is influenced by mountains). To reduce exposure to terminal heat stress while improving yield potential, Gouache et al. (2017) concluded that, under future climate conditions in France, an advancement of stem elongation could be envisaged without increasing frost risk. However, this conclusion must be nuanced at the regional scale. Using ecoclimatic indicators, Le Roux et al. (2024) showed that frost exposure around the 1 cm ear stage will remain a major climatic constraint for wheat production in several French regions, including Limagne. A refined understanding of frost exposure patterns at the (sub-) regional level remains essential for local climate change adaptation, particularly because the spatial drivers of frost risk differ from those governing soil heterogeneity.

Breeding strategies therefore face a dual challenge: reconciling earliness with resilience to low temperatures. Although progress has been made in identifying *loci* controlling phenology and frost tolerance, trade-offs between these traits remain frequent (Babben et al., 2018; Hyles et al., 2020; Soleimani et al., 2022; Zhang et al., 2022). In this context, locally adapted strategies that combine modest acceleration of anthesis with improved cold hardiness may provide robust avenues.

### Limits and perspectives

This study provides a consistent framework to explore how intra-regional soil heterogeneity interacts with climate change in shaping the adaptive value of stress-avoidant wheat ideotypes. However, several simplifying assumptions limit the scope of our conclusions.

We showed the importance of the interaction between elevated [CO_2_] and soil AWC in the determination of the projected impacts of climate change on wheat production and its effects on the identification of stress-avoidant ideotypes. Therefore, the robustness of model predictions critically depends on the formalisms used to represent the effects of elevated [CO_2_] on crop growth and its interactions with soil water availability. Crop models differ in their formalisms to integrate the CO_2_ assimilation of crops under elevated [CO_2_]: they can either represent detailed physiological response (*e*.*g*., biochemical model of leaf photosynthesis including a stomatal conductivity response) or simple empirical relationships (Ewert et al., 2002). In Sirius, the latter approach is used by a applying a linear response to [CO_2_] through the RUE and does not allow any interaction between drought and [CO_2_] *per se*. Therefore, the same proportional yield gains were observed in all soil conditions for a given genotype, but their translation in terms of absolute yield gains differed between soils because of their respective yield potential (Table A.8). This may be oversimplistic as many field studies suggest that the positive effect of [CO_2_] on crop productivity is greater under drought conditions due to direct effect of elevated [CO_2_] on water use efficiency (see the review by Gawinowski et al., 2025). Nevertheless, the severity, the timing and the nature of drought (*i*.*e*., water supply limitation, limited soil water storage, greater evapotranspiration under warmer conditions) will also condition the physiological responses of wheat. For instance, Cao et al. (2022) showed that elevated [CO_2_] modifies the relationships between net photosynthetic rate with the soil water status, either in terms of maximal value or in terms of threshold values. Therefore, the implementation of adequate formalism that allow to robustly account for the impact of elevated [CO_2_] on crop production should be a priority but suffer from a lack of field data to validate crop models under various environments (Gawinowski et al., 2025).

Model initialization relied on assumptions regarding initial conditions and crop management, which may not fully hold under climate change scenario projections. For instance, in the present study, a single sowing date and field capacity at sowing were assumed as initial conditions, whereas future climate conditions may induce delayed or highly variable crop sowing dates due to increased autumn droughts. For example, according to Le Roux et al. (2024), the Limagne plain belongs to an ecoclimatic zone particularly exposed to water deficit in the early stages (from sowing to emergence). Local climate projections suggest a pronounced decline of summer rainfall (Fig. 4) that may deplete soil AWC over the entire soil profile over longer periods. The advantages conferred by deep soils may diminish under prolonged rainfall scarcity, especially if soil recharge does not occur during winter (Wang et al., 2009). Therefore, the robustness of our conclusions should be tested by evaluating the effect of potential shifts of soil water initial conditions, and evaluate their impact on our ideotyping strategy. Similarly, evaluating flexible management options, such as variable sowing dates or densities (*e*.*g*. Flohr et al., 2017, 2020) may complete our vision if, and only if, the anticipation of management changes over time is correct.

## Conclusion

Our study suggests that accounting for local pedological diversity and its interaction with genotype-specific traits may be essential to move from theoretical ideotypes to actionable breeding targets. Soils with limited AWC should not be overlooked, as they often represent a substantial share of cropped areas and are disproportionately affected by the limitations of stress-avoidant strategies. In this case study, the adaptive value of stress-avoidant wheat ideotypes at the regional level appears to be conditional on local soil available water capacity. Deep soils allow earlier stress-avoidant ideotypes to benefit from both terminal stress avoidance and CO_2_ fertilization, whereas shallow soils show more restricted opportunities for these ideotypes.

These findings highlight the need to design ideotypes and adaptation strategies that explicitly integrate pedological diversity. Incorporating high-resolution soil data into crop models and breeding frameworks could be a key step toward regionally relevant adaptation to climate change.

## Funding

The postdoctoral fellowship of G. Blanchet was funded by the International Research Center for Sustainable AgroEcosystems (ISITE CAP20-25).

## Acknowledgments

The authors are grateful to Veronique Genevois and Benjamin Nowak for providing access to local soil datasets and providing guidance on the interpretation of these datasets. We also warmly thank François-Xavier Oury and Alain Chassin for providing local observation data on the Apache cultivar.

## Authors contribution

According to CRediT typology: Conceptualization: G.B., V.A.; Funding acquisition: V.A.; Resources: V.A., M.S.; Investigation: G.B.; Methodology: G.B., M.S., V.A.; Writing—original draft: G.B.; Writing review and editing: G.B., M.S, V.A.; Supervision: V.A.

## Notes

### Competing Interest Statement

The authors have declared no competing interest.

