## Supplementary materials for "How important is the intra-regional soil heterogeneity for the design of future stress-avoidant wheat ideotypes? A modeling study in central France"

^b^ present address: INRAE, UCA, UMR 0547 PIAF, 5 Chemin de Beaulieu, F-63000 Clermont-Ferrand, France

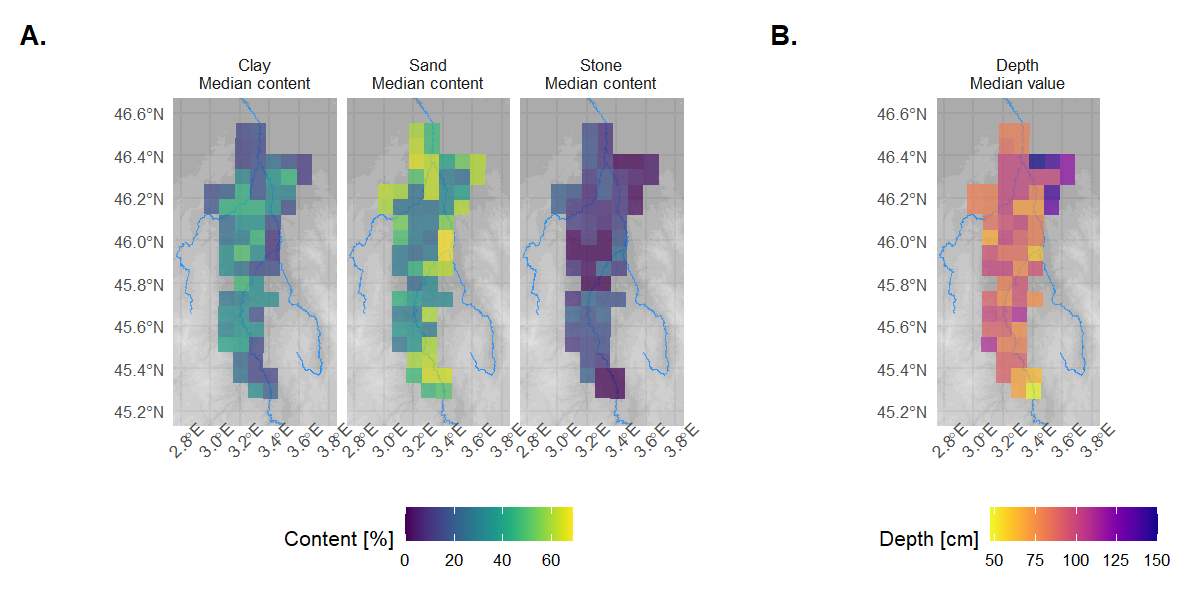

**Fig. A.1.** Spatial distribution of median soil texture (A) and median root-colonizable depth (B) in the Limagne region.

**Table A.2.** **Synthetic validation of LARS-WG generated climate series.** Semi-quantitative summary of LARS-WG diagnostics for the five stations. For each **station** and **variable** (Rain, Tmin, Tmax), **Δμ** is the mean annual bias of **monthly means** (Obs-Gen; units: mm for Rain, °C for Tmin/Tmax)**. Biased months (n/12)** is the number of months with a significant difference in monthly means between observed and generated data (two-sided monthly t-test, p < 0.05). **Global test** reports the p-value class of the paired t-test applied to the 12 monthly means. **Extreme metric** indicates the quantile used to assess extremes, and **Gen/Obs** is the ratio of generated to observed quantiles (P95 for daily precipitation and daily Tmax; P5 for daily Tmin). ns: non-significant; * p < 0.05; ** p < 0.01.

| Variable | Station | Δμ | Biased months (n/12) | Global test | Extreme metric | Gen/Obs |
| --- | --- | --- | --- | --- | --- | --- |
| Rain | MB | -0.02 mm | 0 | ns | P95 | 1.09 |
|  | VC | -2.27 mm | 0 | ns | P95 | 1.16 |
|  | CF | -0.29 mm | 0 | ns | P95 | 1.21 |
|  | IS | 0.57 mm | 0 | ns | P95 | 1.17 |
|  | FT | -1.48 mm | 0 | ns | P95 | 1.11 |
| Tmin | MB | 0.17 °C | 1 | * | P5 | 1.00 |
|  | VC | 0.13 °C | 1 | ns | P5 | 1.00 |
|  | CF | 0.19 °C | 2 | * | P5 | 1.00 |
|  | IS | 0.23 °C | 1 | ** | P5 | 1.00 |
|  | FT | 0.14 °C | 1 | ns | P5 | 1.00 |
| Tmax | MB | −0.19 °C | 2 | ns | P95 | 1.02 |
|  | VC | −0.15 °C | 0 | ns | P95 | 1.02 |
|  | CF | −0.28 °C | 1 | ** | P95 | 1.01 |
|  | IS | -0.03 °C | 0 | ns | P95 | 1.03 |
|  | FT | −0.09 °C | 0 | ns | P95 | 1.02 |

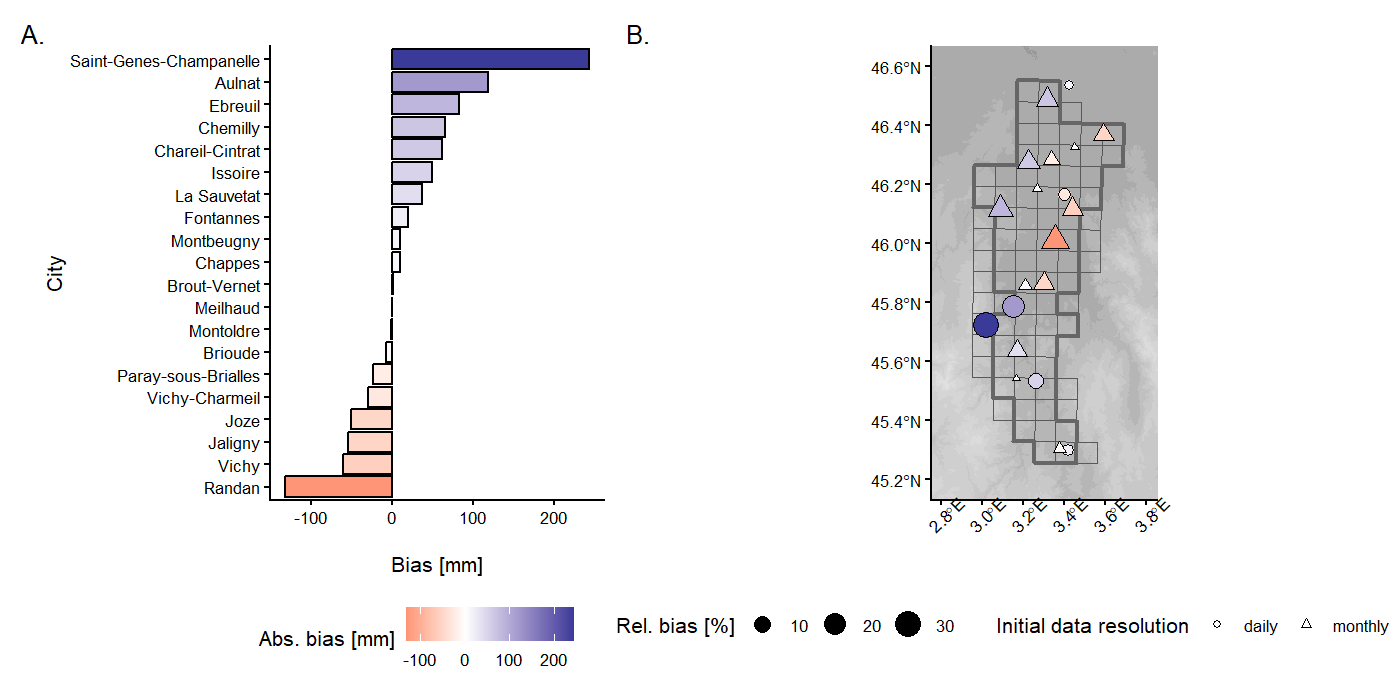

**Fig. A.3.** Evaluation of local annual rainfall bias between Météo-France observations and the SAFRAN interpolated dataset.

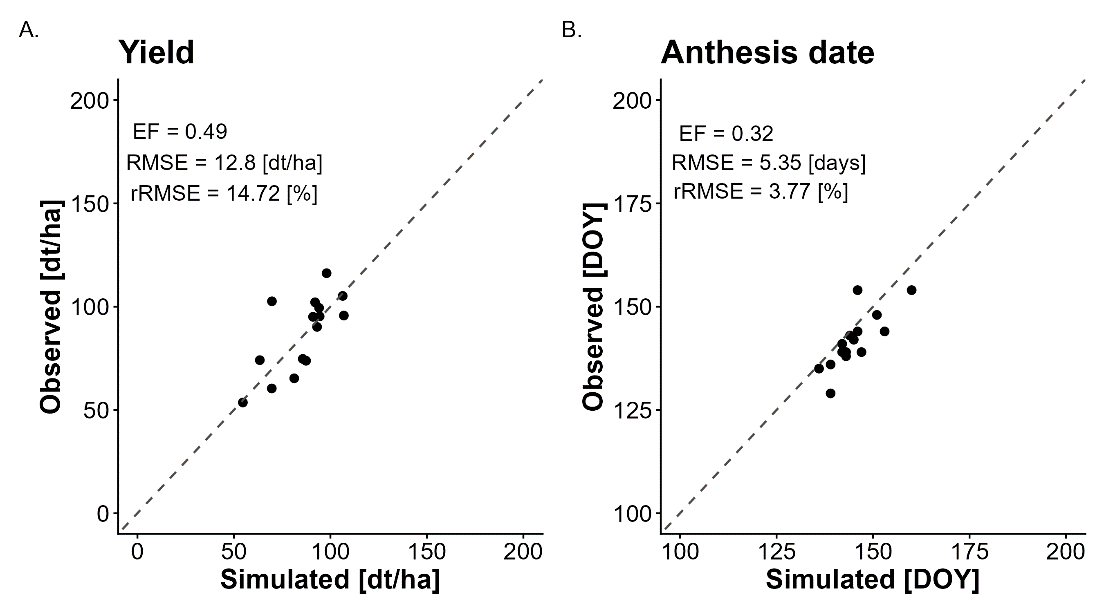

**Fig. A.4**. Evaluation of the Apache cultivar for yield (A.) and anthesis date (B.) prediction at Clermont-Ferrand (16-year observational dataset, 2003–2022).

**Table A.5.** Parameters’ values of the reference genotype (Apache) and parameters’ range explored by the optimization algorithm for generating the virtual genotypes’ panel used in all SSP x GCM x time horizon x site combinations.

| Parameter | Parametrized value of reference genotype | Parameter range for virtual genotypes’ panel | | References |
| --- | --- | --- | --- | --- |
|  |  | Lower bound | Higher bound |  |
| PHYLL | 100.6 | - | - |  |
| VAI | 0.0014 | 0.0004 | 0.0022 | Robertson et al. (1996),  He et al. (2012), |
| VBEE | 0.022 | 0.002 | 0.202 | Robertson et al (1996).  He et al. (2012), |
| SLDL | 0.639 | 0.05 | 0.957 | He et al. (2012), |
| DSGNR_max_ | 0.237 | - | - |  |
| DSGNS | 0.199 | - | - |  |
| DSGNT | 0.9 | - | - |  |

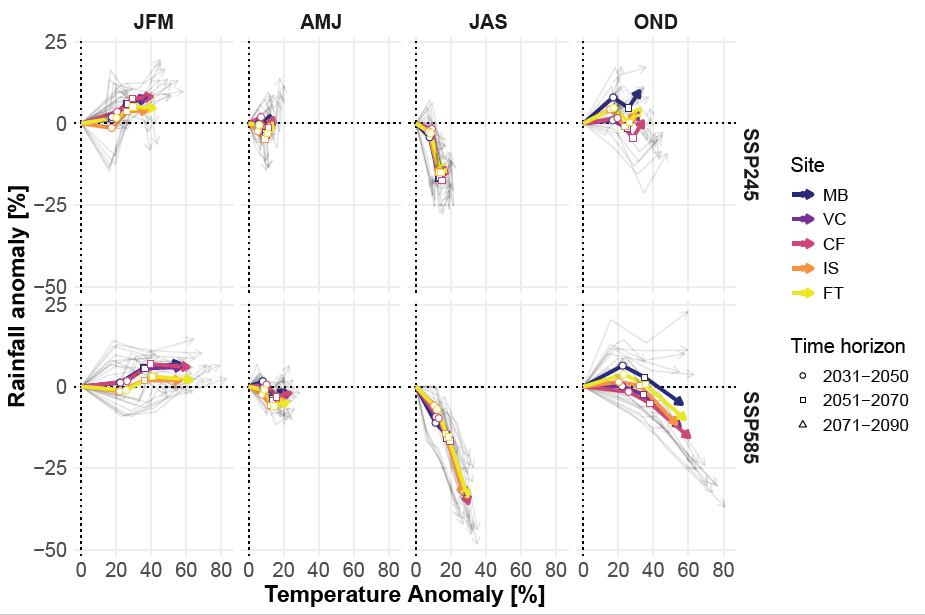

**Fig. A.6.** Seasonal trajectories of absolute temperature and precipitation anomalies for each site. Anomalies were calculated relative to the baseline period (1990-2020). Each arrow node corresponds to the coordinates of the anomalies at a given time horizon. Thin gray arrows show the trajectories for each site-GCM combination while thick-colored arrows show the median trajectories for each site.

**Table A.7**. Mean wheat yield (dt ha⁻¹) across soils × sites × GCMs by SSP scenario and time horizon, with vs without CO₂ elevation. Values are means (SD).

| **Time period** |  | **2031-2050** | **2051-2070** | **2071-2090** |
| --- | --- | --- | --- | --- |
| **SSP scenario** | **[CO_2_] elevation** |  |  |  |
| 2-4.5 | No | 57.9 (24.0) | 56.9 (23.8) | 58.7 (24.4) |
|  | Yes | 62.6 (26.0) | 64.6 (27.1) | 68.9 (28.7) |
| 5-8.5 | No | 57.6 (24.1) | 55.8 (24.0) | 55.9 (23.7) |
|  | Yes | 63.5 (26.7) | 68.7 (29.6) | 73.7 (31.5) |

**Table A.8.** Mean CO₂-driven yield increase (dt.ha^-1^) across all sites × GCMs by soil depth category, climate scenario and time horizon. Values are means (SD).

| **Time period** |  | **2031-2050** | **2051-2070** | **2071-2090** |
| --- | --- | --- | --- | --- |
| **SSP scenario** | **Soil depth** |  |  |  |
| 2-4.5 | Shallow | +3.96 (1.90) | +6.47 (2.96) | +8.62 (3.89) |
|  | Deep | +5.59 (2.19) | +9.18 (3.46) | +12.3 (4.46) |
| 5-8.5 | Shallow | +5.05 (2.39) | +10.7 (4.89) | +14.8 (6.65) |
|  | Deep | +7.12 (2.77) | +15.6 (5.80) | +21.7 (7.80) |
